# Topology-Dependent Interference of Circuit Function by Growth Feedback

**DOI:** 10.1101/724906

**Authors:** Rong Zhang, Jiao Li, Juan Melendez-Alvarez, Xingwen Chen, Patrick Sochor, Qi Zhang, Hanah Goetz, Tian Ding, Xiao Wang, Xiao-Jun Tian

## Abstract

Growth-mediated feedback between synthetic gene circuits and host organisms leads to diverse emerged behaviors, including growth bistability and enhanced ultrasensitivity. However, the range of possible impacts of growth feedback on different gene circuits remains underexplored. Here, we mathematically and experimentally demonstrated that growth feedback affects the functions of memory gene circuits in a network topology-dependent way. Specifically, the memory of the self-activation circuit is quickly lost due to the fast growth-mediated dilution of the circuit products. Decoupling of growth feedback reveals its memory, manifested by its hysteresis property across a broad range of stimulus. On the contrary, the toggle switch is more refractory to the growth-mediated dilution and can retrieve its memory after the fast-growth phase. The underlying principle lies in the different dependence of active and repressive regulations in these circuits on the growth-mediated dilution. Our results unveil the topology-dependent mechanism on how growth-mediated feedback influences the behaviors of gene circuits.

## INTRODUCTION

Circuit-host interactions affect behaviors of synthetic gene circuits, thereby adding an additional layer of complexity to already intricate regulation networks^1–3^. There are many sources of circuit-host interactions, including metabolic burden^4,5^, cell growth^6–10^, and resource relocation/competition^11–14^. These interactions are often neglected in the design of gene circuits based on the assumption that gene circuits are generally orthogonal to the host background^15–18^. In many instances, however, the impact of interactions between the gene circuit and the host organism is significant^19–21^. Understanding the mechanisms of how circuit-host mutual interactions are established, particularly the effects of these interactions on gene circuit functions will help us to better formulate control strategies for designing and engineering robust gene circuits that ensure improved accuracy in applications.

Various feedback loops are created by circuit-host interaction. For example, we know that growth feedback is formed based on the fact that the expression of the synthetic gene circuits inevitably causes metabolic burden to the host cells and thus affects cell growth, which in turn changes the gene expression of circuits^22,23^. This growth-mediated feedback can endow synthetic gene circuits with various emerged properties. An example of this would be of enhanced ultrasensitivity by growth feedback, which makes a non-cooperative positive autoregulatory circuit bistable^24,25^. Another example is the innate growth bistability occurrence in drug-resistant bacteria when an antibiotic resistance gene expression is coupled to cell growth^26^. Toxin cooperativity can also be induced in the multiple toxin-antitoxin systems with several growth-mediated feedbacks^27^. However, the desired function of gene circuits can also be attenuated or disabled by the growth-mediated feedback and thus can be evaluated inaccurately.

In this paper, we present a study of how network topology affects the extent to which the growth feedback influences gene circuit behaviors. We used several synthetic memory gene circuits with different topologies to test whether the circuit-host interaction can affect the memory maintenance function of these circuits. We know in the literature of several studies on synthetic switch circuits, including the toggle switch, self-activation switch, push-on-push-off switch, quorum-sensing switch, and quadrastable switch^13,28–32^. To test the hysteresis properties of these circuits, a dilution protocol was often used to ensure the cells in the exponential growth phase. For example, in the seminal work on the toggle switch, all the samples were washed and diluted in fresh medium every 6-10 hours^28^. The underlying reason for this is that cell growth can be maintained at a relatively constant rate. While this dilution protocol works successfully to demonstrate the hysteresis properties of the circuits, an important question is whether constant cell growth is a necessary condition for these circuits to show the bistability and whether bistable behavior can also be observed robustly in conditions where the growth rate is rather dynamic instead.

In our experiments, we systematically tested the dynamic behaviors of the self-activation switch and toggle switch under dynamic cell growth condition. We found that the memory of the self-activation switch is lost due to cell growth and thus no hysteresis is found using the dilution protocol with fresh medium. However, after uncoupling the growth feedback from gene circuits using the protocols of limiting cell growth, we found a broad range of bistability. We further tested the toggle switch with our protocols and found that it is more refractory to growth feedback. Thus, we concluded that the effects of growth-mediated feedback on gene circuits depend on their network topologies. While some circuits are robust to the perturbation from growth feedback, the functions of others are easily compromised. We validated this underlying mechanism using mathematical modeling and theoretical analysis by integrating the dynamics of cell growth into the gene circuits.

## RESULTS

### Theoretical analysis of the self-activation circuit reveals bistability, but experimental data shows no memory

To determine the effects of growth feedback on the gene circuits, we first built a simple self-activation gene circuit, in which the transcriptional factor AraC forms a dimer and binds to the promoter P_BAD_ in the presence of stimulus arabinose (L-ara), which in turn activates the expression of itself and the reporter GFP. We measured the dose-response curve of the prompter P_BAD_, which shows ultrasensitivity after parameter fitting (Fig. 1A). Based on the fitted parameters, we developed a mathematical model for the AraC self-activation system (see Methods for details). Through analysis, we predicted that this system is bistable under the right conditions, i.e., as shown in Fig. 1B, the system suddenly switches on at a given threshold as the stimulus increases if the initial state is ‘OFF’. Once activated, the system stays on the ‘ON’ state even if the stimulus is completely removed. A theoretical analysis of our mathematical model, parameterized with experimental data, predicts that the AraC self-activation circuit is bistable.

**Figure 1.**
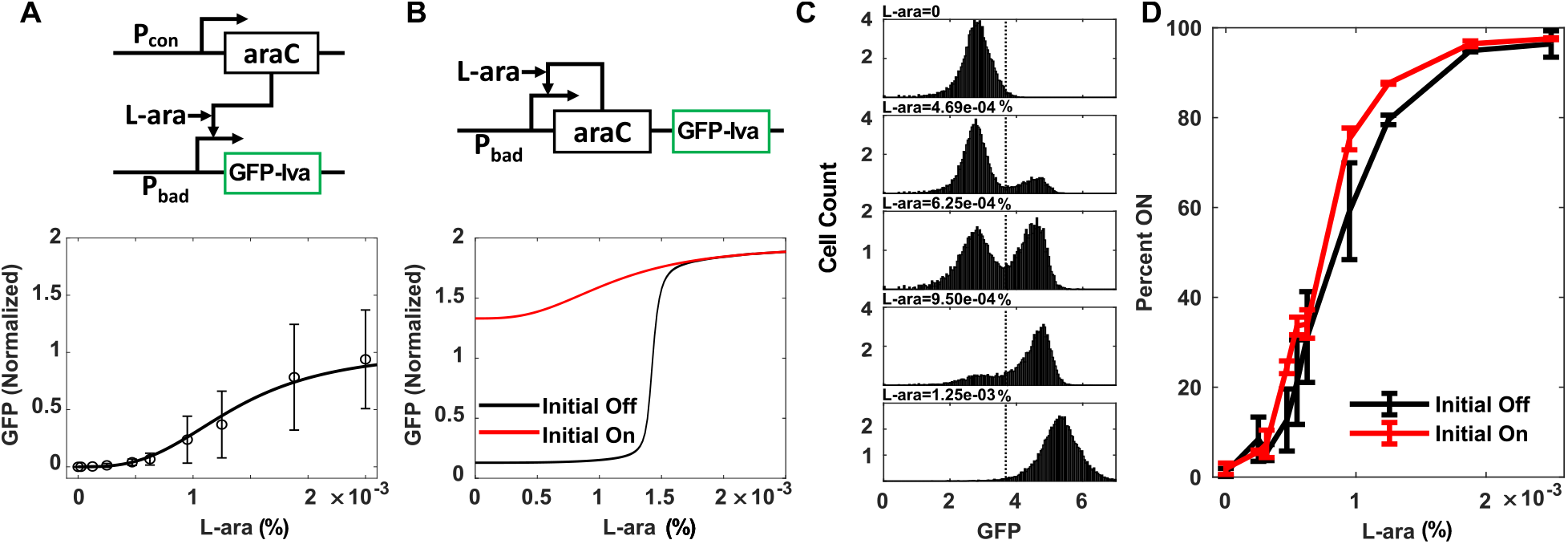
Theoretical analysis reveals bistability, but experimental data shows no hysteresis. **(A)** Parameter fitting the dose-response curve of the prompter (P_BAD_) shows ultrasensitivity. The results showed the mean value of 6 replicates. Mean of normalized value was shown. Unstable GFP variant (GFP-lva) is used as the reporter. **(B)** Mathematical model predicts bistability from AraC self-activation circuit with P_BAD_ prompter. **(C)** Steady-state distribution of the GFP after 17 hours induction of inducer L-ara with the initial state “OFF” shows two distinct states (separated by the dash lines), ‘ON’ and ‘OFF’. **(D)** The steady-state fraction of ‘ON’ cell after 17 hours induction of inducer L-ara with the initial state “OFF” (blue curve) or ‘ON’ (Red curve) shows no hysteresis. Data displayed as mean value of three replicates.

To further test our prediction, we measured the response of AraC self-activation circuit to varying doses of stimulus L-ara. Clear bimodal distributions were observed in a broad range of arabinose concentrations (Fig. 1C), with some of the cells in the high GFP state and some in the low GFP state, suggesting the system could be bistable. To verify the bistability, further experiments were carried out to test for hysteresis. The fraction of ‘ON’ cells was measured by flow cytometry under two different initial conditions and was plotted as a function of arabinose concentration (Fig. 1D). First, cells were initially in the inactive state and then treated with a series dose of L-ara for 17 hours. The fractions of activated cells established a threshold above which the system can be fully activated (Fig. 1D, black line). Second, the activated cells pretreated with a high dose (2.5E-3%) of L-ara were washed twice with fresh LB, then diluted 100 times into fresh LB with various concentrations of arabinose. After 18 hours of culturing, the fraction of activated cells showed a curve similar to the one without pretreatment (Fig. 1D, red line), which was confirmed by the similar steady-state GFP curves measured by plate reader with different initial states (Supplementary Fig. 1). These data suggest that this self-activation circuit is either bistable within a tiny stimulus range or monostable. Alternatively, the system is actually bistable in a broad range of stimulus but the memory of circuits within the activated cells was lost due to some additional implicit links involved in the system. Thus, seemingly inconsistent conclusions are found from theoretical prediction and experimental verification.

### Growth-mediated feedback disguises the bistability of the self-activation circuit

To uncover the underlying reason for the memory loss, we further studied the temporal dynamics of activated cells post-dilution in fresh medium. After diluting the activated cells with 1:100 ratios in fresh media with various concentration of L-ara, the cell density (measured as optical density (OD) at 600 nm) and GFP fluorescence were measured at different time points. We observed that cell growth rate for all the samples decreased as the population reaching the carrying capacity, following the logistic model? (Fig. 2A). However, the average GFP level (GFP/OD) decreased rapidly within the exponential growth phase and reached a very low level at 3 hours after dilution (Fig. 2B), indicating the memory loss of the circuit. Later on, GFP started accumulating and then reached different steady states depending on the L-ara levels (Fig. 2B), which generated curves similar to the ones started from the inactive state (Supplementary Fig. 2). It is important to note that the GFP level maintained very well for 6 hours when the cells reached the stationary phase, although a small decrease was found at around 10 hours. To further confirm that memory loss did not result from the factors associated with the stationary phase, the activated cells in the exponential phase were diluted into fresh medium. As shown in Supplementary Fig. 3, cells were activated by a high dose of L-ara to different levels at OD=0.3~0.7 and still all of them showed a decline of GFP after diluting into fresh medium, consistent with the results of the dilution of the stationary phase cells into fresh medium. Similar phenomena were also found in the LuxR self-activation circuit (Supplementary Fig. 4). Thus, fast cell division indeed can dilute the synthetic circuit’s gene expression.

**Figure 2.**
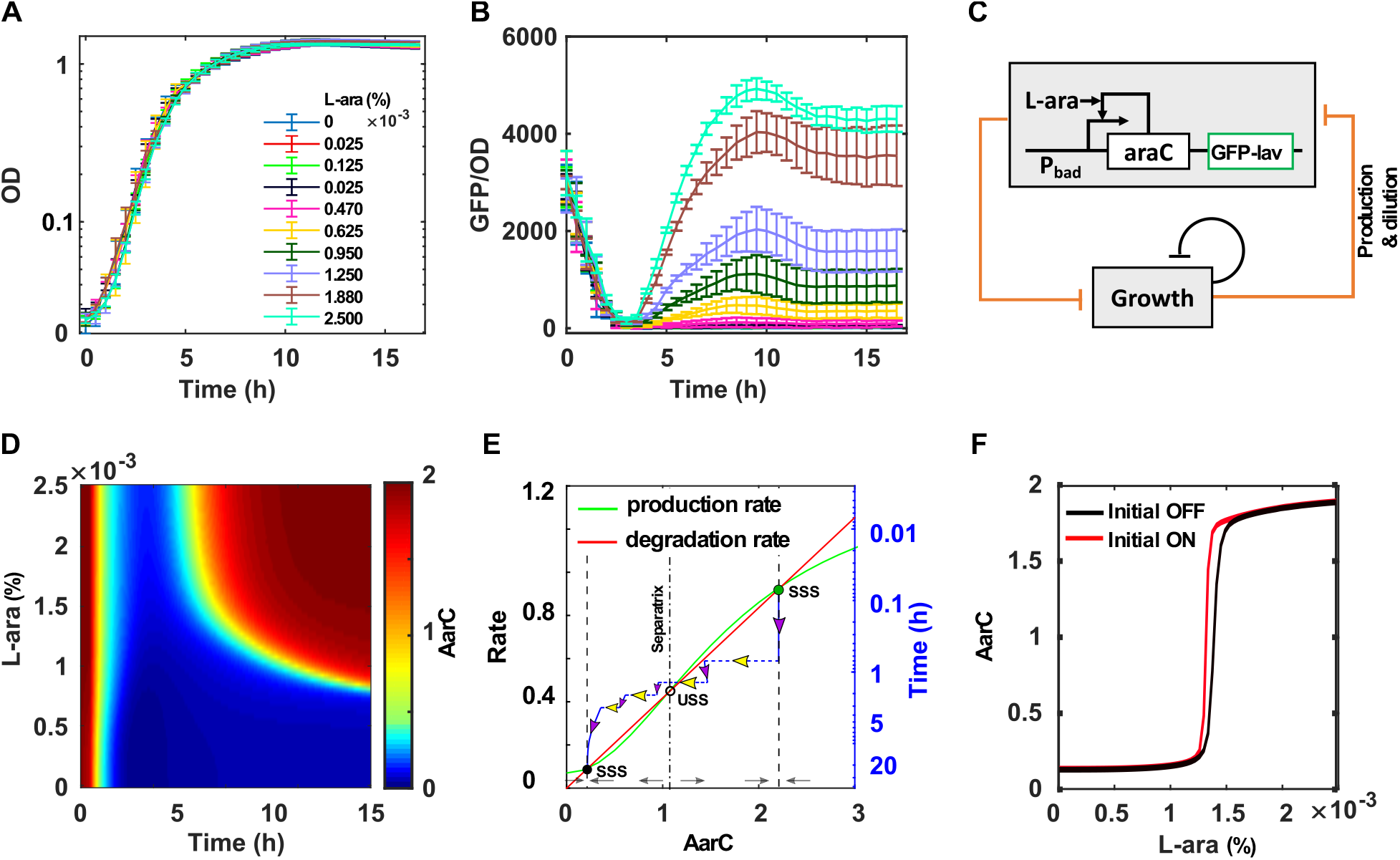
Growth-mediated feedback disguises the bistability of self-activation circuits. **(A-B)** Dynamics of growth (Optical Density (OD)) (A) and GFP/OD (B) after 1:100 dilution of ‘ON’ cells into the fresh medium with various concentrations of L-ara. Mean value of three replicates were shown. **(C)** Diagram of coupling gene circuit with cell growth. The expression of gene circuit causes a burden to the cell and slow down; the cell growth in turn dilutes or decreases the expression of the gene circuit, forming growth feedback. The self-inhibition of the cell growth indicates growth rate decreases as cell density increases. The mathematical model is revised based on this diagram. **(D)** Simulation with revised mathematical model shows the change of AraC as a function of time and dose of inducer L-ara. The system was set to the ‘ON’ state initially. **(E)** The process of memory loss of the self-activation switch. Simulated trajectory (blue lines) of one cell is shown on the rate-balance plot of AraC. Two stable steady states (SSS, solid circles) and one unstable steady state (USS, open circle) are shown at the intersection of production rate curve (green) and degradation curve (red). The system was set to the ‘ON’ state initially (solid green circle) and L-ara was set to 0 to test the memory maintenance. Four cell division events were considered in simulation (dashed line with yellow arrow). The separatrix (dash-dotted line) determines whether the system goes to the ‘ON’ state or the ‘OFF’ state as shown by the grew arrows. The rate-balance plot is based on the model without growth feedback. **(F)** Simulation confirms that the bistable range of self-activation circuit is significantly reduced when coupled with growth feedback compared with Fig.1B.

We also found that the growth rate decreased for the cells that contain the synthetic circuit compared with cells without any circuit (Supplementary Fig. 5), which is consistent with previous findings that synthetic circuits cause burdens to the cells and inhibit cell growth^7,23,24,33^. All these data suggest that dilution with fresh medium brings a growth-mediated feedback loop into the synthetic gene circuit, in which expression of the gene circuit slows down cell growth, whereas fast cell growth dilutes the expression of gene circuits (Fig. 2C). That is, once the activated cells were diluted into fresh medium, there was a quick increase of cell growth that significantly diluted the gene expression of the synthetic circuits, leading to the decrease of gene products and thus inactivation of the circuit.

To further understand how cell growth changes circuit behavior, we revised our mathematical model by integrating the growth-mediated feedback (see Methods for details). We conducted a simulation to demonstrate how the AraC self-activation circuit lost memory after diluting into the fresh medium at both populational and single-cell levels. At the populational level, a full spectrum of AraC dynamics under different levels of L-ara is shown in Fig. 2D. Initially, AraC was set at a high level but decreased very quickly to low levels due to dilution from fast cell growth, consistent with the experimental results (Fig. 2B). At the single-cell level, we developed an algorithm to simulate the interplay between the circuit and cell growth by considering the stochasticity from cell division before the system reaches the stationary phase (See Method for details). Four cell division events were considered for one cell between post-dilution and pre-stationary phase. As shown in Fig. 2E, two stable steady states (SSS) and one unstable steady state (USS) can be found at the intersections of the production and degradation rates of AraC (green and red curves respectively). The AraC level was set to high level at the ‘ON’ state (green circle in Fig. 2E) initially, and then decreased because of cell division-mediated dilution (dashed lines with yellow arrows in Fig. 2E), and later started accumulating (solid blue lines with purple arrows in Fig. 2E) because of Ara C production by the self-activation circuit. However, the dilution was much faster than the accumulation, and thus AraC levels decreased very quickly to a point below the separatrix (dash-dotted line in Fig. 2E) and converged to the low level at the ‘OFF’ state (black circle in Fig. 2E), and maintained in this state even after the cell growth became slower in the pre-stationary and stops at the stationary phases. That is, at the early stage, the growth feedback was dominant and continuously diluted the circuit expression, leading to switch-off of the circuit. This is the underlying mechanism for the memory loss of the AraC self-activation circuit and the reason why no hysteresis was found using the dilution protocol with fresh media (Fig. 1D, Fig. 2F).

For theoretical analysis, we also considered a scenario of constant growth rate to study how the strength of growth feedback affects the gene circuits (see Methods for details). Increase the growth rate indeed changes the bifurcation diagram significantly (Supplementary Fig. 6A). The activation threshold of the switch increases exponentially with the growth rate (Supplementary Fig. 6B). Furthermore, the deactivation threshold can be changed from negative to positive, making the switch reversible instead of irreversible (Supplementary Fig. 6). This suggests that it is unlikely to activate the system in the experiment under fast-growth conditions. It is worth noting that the stimulus range for bistability is expanded with growth rate, consistent with previous findings that growth feedback increases the nonlinearity of one non-ultrasensitive self-activation circuit and makes it bistable^24^. Taken together, these data suggest that the underlying mechanism for the memory loss of the self-activation circuit lies in the growth feedback. That is, growth-mediated feedback disguises the bistable property of the self-activation circuit.

### Decoupling of growth feedback reveals the bistability of the self-activation circuit

The above results showed that dilution with the fresh medium can keep cells fast-growing, but inevitably it induces the growth feedback to synthetic gene circuits, leading to memory loss. To decouple the growth feedback from the circuits and overcome this adverse effect, we developed a new dilution protocol. Instead of dilution into the fresh medium, we diluted cells into the conditioned medium that maintains the cells in the stationary phase. The conditioned medium was collected from the samples that were cultured without any induction in parallel (Fig. 3A, left panel). The growth feedback was expected to be decoupled (Fig. 3A, right panel).

**Figure 3.**
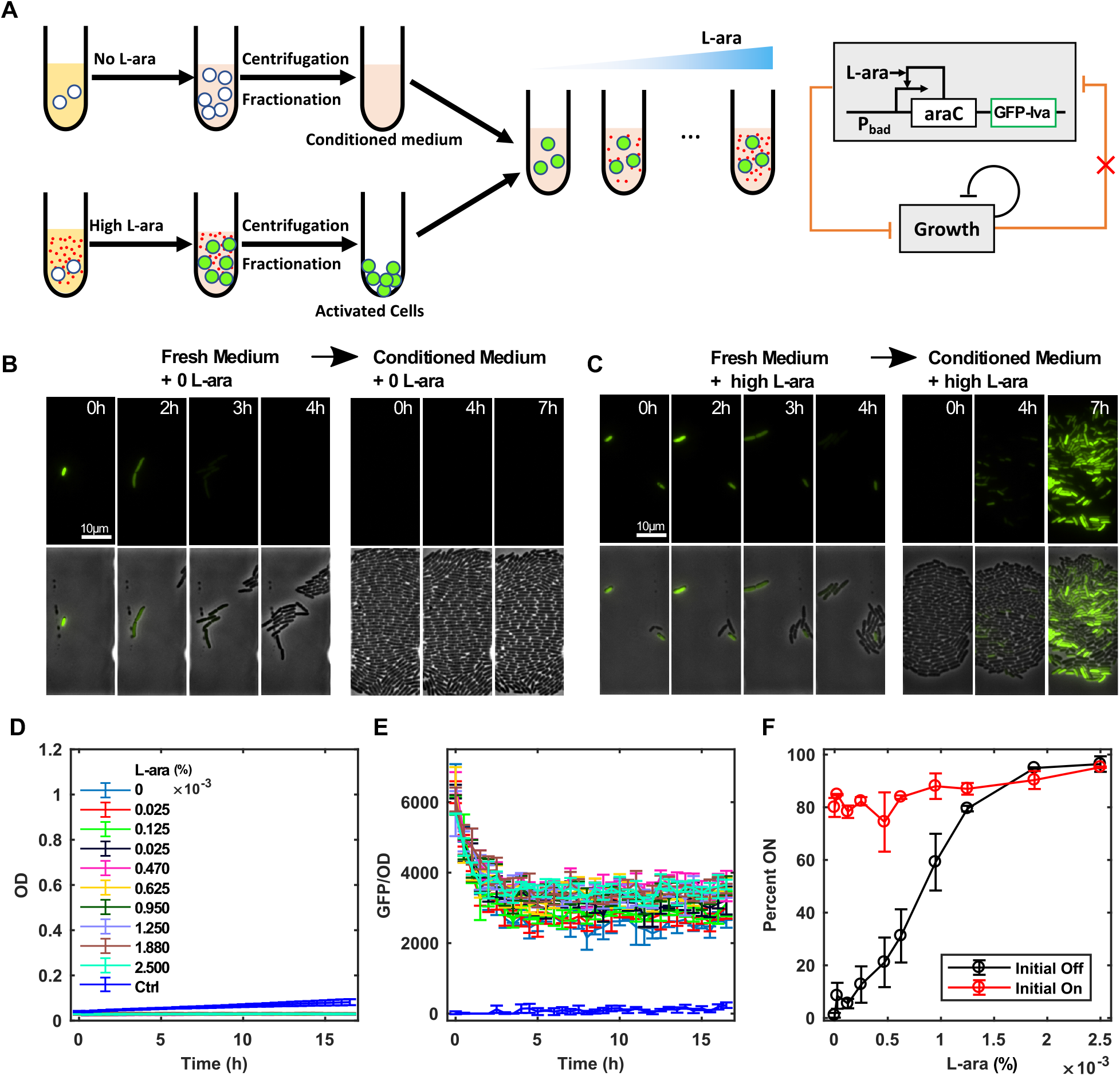
Decoupling of growth feedback reveals the bistability of the self-activation circuit. **(A)** Diagram of the new protocol decoupling of the growth feedback. Instead of dilution into fresh medium, cells were diluted into conditioned medium, which is from a parallel culture without any inducers in the stationary phase. **(B-C)** Time-lapse imaging of GFP (and brightfield overlay) in the AraC self-activation circuit showed that it switches off after several rounds of cell divisions with fast growth in fresh medium and then did not recover in the conditioned medium without inducer (B) but recovered with a high dose L-ara (C). Representative results from three replicates are shown. **(D-E)** The dynamics of growth (D) and GFP (E) after 1:100 dilution of activated cells into the conditioned medium with various concentration of L-ara. Negative control (blue line) was ‘OFF’ cells grown in conditioned medium with no L-ara. Data indicate the mean±SD of three independent replicates. **(F)** Hysteresis curves obtained using the new protocol by diluting ‘ON’ cells into the conditioned medium with various concentrations of L-ara. Mean value of three replicate tests were shown.

To test this protocol, the activated cells were loaded into a microfluidic platform and fresh medium with or without L-ara was provided at first. We found that the GFP level was extinguished quickly along with several cell divisions regardless the L-ara levels (Fig. 3B-C and Supplementary Video 1-2), thereby confirming the memory loss of the self-activation switch due to the growth feedback (Fig. 2B). However, after switching to conditioned medium, we observed that the GFP level did not recover without L-ara (Fig. 3B and Supplementary Video 1) but it elevated under a high dose of L-ara (Fig. 3C and Supplementary Video 2), which is consistent with the results in Fig. 2B. Furthermore, we measured the dynamics of the systems under the conditioned medium at the populational level. We found that the cells did not divide as the OD was maintained in the low level (Fig. 3D) but still had a strong ability to express the genes of the circuit since the GFP levels were maintained very well during 17 hours for all L-ara concentrations (Fig. 3E). It is noted that the maintenance of GFP is not from slow degradation as the unstable GFP variant (GFP-lva) with a half-time of 40min was used in the circuit^34^. This is consistent with the finding that a surprisingly long period of constant protein production activity has been observed in the stationary phase of bacteria [19]. Finally, a very large hysteresis range from the protocol with the conditioned medium was revealed. As shown in Fig. 3F, the cells were pretreated with a high dose of inducer L-ara and then diluted into the conditioned medium with various concentrations of inducer. The fraction of the activated cells showed strong stability, indicating that the system actually functions as an irreversible bistable switch, consistent with the prediction from our mathematical model (Fig. 1B). These data suggest that growth feedback can be decoupled with conditioned medium. To further confirm this conclusion, we diluted the activated cells into low nutrition medium M9 with either no L-ara or a high dose of L-ara. The memory was maintained very well in both cases of low nutrition medium (Supplementary Fig. 7A), in contrast with the samples diluted in high nutrition medium (Supplementary Fig. 7B). Taken together, decoupling the growth feedback with conditioned medium from the stationary phase or low-nutrient medium helps maintain the memory of the gene circuits and reveals the bistability of the self-activation circuit.

### Toggle switch is refractory to memory loss resulted from the growth-mediated feedback

We have tested the effect of the growth-mediated feedback on the synthetic self-activation circuit and found that growth-mediated feedback disguised the bistable behavior of this circuit. In order to study whether the growth feedback also affects other types of gene circuits, we tested our protocol with the toggle switch, in which two genes tetR and LacI inhibit each other. In the original work, the authors used a method to dilute cells into the fresh medium frequently in order to maintain the cell in the log growth phase^28^. The hysteresis behavior of the toggle switch was indeed found. However, the effects of growth-mediated feedback on the toggle switch are not clear. Thus, we studied the dynamics of the toggle switch with our protocols. At the populational level, after dilution of activated cells in fresh medium, which couples the growth feedback with the toggle switch circuit (Fig. 4A), the GFP was seen to decrease with time to the minimum at 4 hours after dilution before reaching the maximum levels at 16 hours (Fig. 4B-C). However, it is worthwhile to note that the GFP expression was not completely switched off and was reactivated for all the samples with various concentrations of inducer aTc, indicating that the memory of the toggle switch was maintained during the fast-growth phase and can be retrieved during the slow-growth phase. Similarly, after dilution of activated cells into conditioned medium to decouple the growth feedback (Fig. 4D), GFP levels of the toggle switch were also maintained very well for 17 hours (Fig. 4E-F). It is noted that the maintenance of GFP is not from slow degradation as the unstable GFP variant (GFP-laa), which has the same half-time as GFP-lva, was used in the circuit^34,35^.

**Figure 4.**
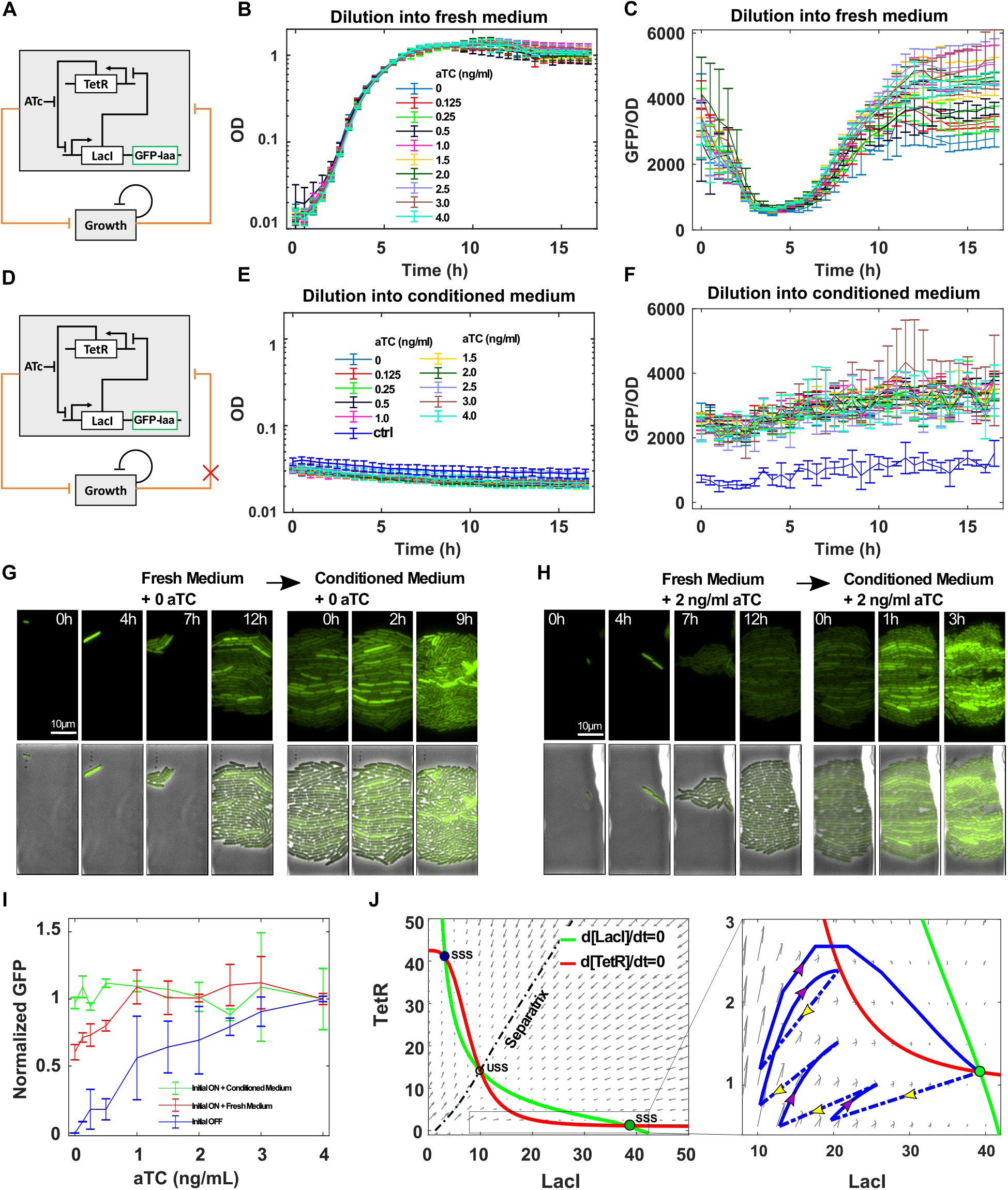
Toggle switch is refractory to memory loss from the growth-mediated feedback. **(A)** Diagram of the toggle switch circuit coupled with growth feedback in fresh medium. Unstable GFP variant (GFP-laa) is used as the reporter. **(B-C)** Dynamics of cell growth and GFP/OD after 1:100 dilution of ‘ON’ cells into fresh medium with various concentrations of inducer aTc. Data indicates the mean±SD of three independent replicates. **(D)** Diagram of the toggle switch circuit decoupled with growth feedback in condition medium. **(E-F)** Dynamics of OD and GFP/OD after 1:100 dilution of ‘ON’ cells into the conditioned medium with various concentration of inducer aTc. Negative control (blue line) was ‘OFF’ cells grown in conditioned medium without aTc. Data indicate the mean±SD of three independent replicates. **(G-H)** Time-lapse imaging of GFP (and brightfield overlay) in the toggle switch circuit showed that it decreased after several rounds of cell divisions with fast growth in fresh medium and then recovered in the conditioned medium under both the condition without inducer (G) or with a high dose of inducer ATc (H). Representative results from three replicates are shown. **(I)** Hysteresis curves generated by dilution of ‘ON’ cells either into conditioned medium (green curve) or fresh medium (red curve) with different concentrations of aTc. Three replicate tests were performed to generate each curve. The GFP level was normalized to the level at the highest aTc concentration. Mean±SD of the normalized value of each curve was shown. **(J)** Simulated trajectory (blue lines) of one cell is shown in the direction field of LacI/TetR. Four cell division events were considered (indicated by dashed lines with yellow arrows in the enlarged box area). Two stable steady states (SSS, solid circles) and one unstable steady state (USS, open circle) are shown at the intersection of nullclines (green and red curves). The system was set to the ‘ON’ state initially (solid green circle) and aTc was set to 0 to test the memory maintenance. Four cell division events were considered in simulation (dashed line with yellow arrow). The separatrix (dash-dotted line) determines whether the system goes to the ‘ON’ state or the ‘OFF’ state as shown by the grew arrows. The nullcline curves were based on the model without growth feedback.

At the single-cell level with microfluidics, the GFP level decreased with cell division in the fresh medium regardless of the concentrations of inducer aTc as well, and the system was reactivated after switching to the conditioned medium both with or without inducer aTc (Fig. 4G-H and Supplementary Video 3-4). Although GFP expression showed heterogeneity among cells (Fig. 4G-H), it is noted that it did not completely switch off, which is consistent with Fig. 4C. Furthermore, the system needed more time to recover without inducer aTc (Fig. 4G and Supplementary Video 3) in contrast to the high dose (Fig. 4H and Supplementary Video 4). In Fig. 4I, the hysteresis curves from conditioned medium (green curve) and the fresh medium (red curve) both showed the irreversibility of the toggle switch, consistent with the previous works^28,36,37^. Thus, the toggle switch is refractory to memory loss from the growth-mediated feedback.

To demonstrate the underlying mechanism of how the differences in the topology differ the response of the self-activation switch and toggle switch to growth feedback, simulations were done to analyze the behavior of the toggle switch. First, at the populational level, no difference was found in the hysteresis curves for fast growth and no growth conditions (Supplementary Fig. 8), consistent with experimental results (Fig. 4I). At the single-cell level, the trajectory of one cell is shown to demonstrate how the system changes with several cell divisions and then recovers to the original state. As shown in the directed field for the toggle switch (Fig. 4J), two stable steady states (SSS) and one unstable steady states (USS) can be found at the intersections of the nullclines. The cell was set initially at the High-LacI/Low-TetR state (‘On’ state, solid green circle in Fig. 4J) and then moved toward the left-bottom corner (Fig. 4J, blue dashed lines with yellow arrows) due to the cell division. It tried to recover toward the right-top corner (Fig. 4J, solid blue lines with purple arrows) after each cell division. After several rounds of competition between the accumulation and growth-mediated dilution, the system reached a stationary growth phase in which cells dit not divide anymore. Here, the system was still in the domain where it can easily recover to the original High-LacI/Low-TetR state.

The major differences between the self-activation switch and the toggle switch are the promoters of the genes used in the circuits. The promoter p_BAD_ in the self-activation circuit is positively regulated by the transcriptional factor AraC, while the promoter p_Tet_ in the toggle switch is negatively regulated by the transcriptional factor tertR. Given that the activity of the inducible promoter p_BAD_ directly depends on the concentration of the transcription activator AraC, the dilution of the activator significantly decreases the production rate of the gene in the self-activation switch. While the activity of the inducible promoter p_Tet_ inversely depends on the concentration of the transcription repressor tetR, the dilution of the repressor does not decrease the relative production rate of the two mutual-inhibitive genes and thus leads to the robust memory (Supplementary Fig. 9). These differences between the toggle switch and the self-activation switch are also reflected in the directions of the dilution (dashed lines with the yellow arrows in Fig. 2E and Fig. 4J). The dilution direction is more parallel to the separatrix for the toggle switch (dash-dotted line in Fig. 4J), while it is perpendicular to the separatrix for the self-activation switch (dash-dotted line in Fig. 2E). That is the reason why the system has crossed the separatrix after two cell divisions for the self-activation circuit, while it still has not crossed the separatrix after four cell divisions for the toggle switch. In summary, the network topology of the toggle switch makes it robust to the growth-mediated feedback.

## DISCUSSION

### Host-mediated interference of circuit function depends on the topologies of the synthetic gene circuit

It is still a big challenge to build large-sized synthetic gene circuits since oftentimes circuits do not function as expected once they are assembled. One fundamental reason is that multiple factors of circuit-host interactions, such as metabolic burden, cell growth feedback, and resource competition are often neglected when building and testing gene circuits because the assumption is that these circuits are orthogonal to the host background. However, we know that physiological links from the host to the gene circuits create a hidden regulatory layer for synthetic gene circuits, which often perturbs the expected functions of gene circuits. In addition, it is difficult to predict how these hidden interactions affect the circuit function and how we can minimize the unfavorable effects. Our data suggest that the circuit-host interaction mediated by growth feedback may disguise the true behavior of the synthetic gene circuits. Most importantly, the interference depends on the network topology of the gene circuit. While the self-activation circuit is very sensitive to the growth feedback and loses its memory easily, the toggle switch is robust to the growth feedback and can maintain the circuit’s memory very well even though some decline has been found due to fast cell growth.

To decouple the growth feedback from the gene circuits, we tested protocols with the conditioned medium from the stationary phase or the low-nutrient medium that stops or slows down the cell growth. We found that the effects of the growth-mediated feedback can be minimized and the true behavior of the gene circuit finally exhibited. This is consistent with the finding of different dependences on simple negative and positive regulation on growth rate^23^. The expression of one gene increases with the increase of growth rate for a negative regulation but decreases for a positive regulation^23^. Given that most synthetic gene circuits are composed of simple negative and positive regulations, the circuit-host interactions affect each link differently and thus can significantly change the functions of the gene circuits. Furthermore, it will be interesting to study how the growth-related changes in global gene expression noise affect the behavior of the circuits given that the noise level is coupled with the growth rate^38^. The analysis of input-associated Signed Activation Time (iSAT) can be used in the design of robust switches that maintain a reliable memory^39,40^.

### Circuit-host mutual interactions hold a double-edged effect on the gene circuits

Here, we found that growth feedback has an adverse effect on the functions of some gene circuits as the memory can be lost due to fast cell growth. Previously, it has also been found that the growth feedback lends some new properties to some gene circuits. A constitutively expressed antibiotic resistance gene, for example, shows innate growth bistability given that the expression of the resistance gene is growth-dependent and the cell growth is modulated by translation-targeting antibiotics^26^. Increased cell growth increases the expression of the resistance gene, which in turn reduces the antibiotic concentration in the cell and further promotes cell growth. It is also reported that growth feedback makes some other circuit feedback systems more cooperative^23,24^. A non-cooperative circuit, autoregulation of T7 RNA polymerase, was found to generate bistability unexpectedly^24^. The underlying mechanism is that the nonlinear dilution of T7 RNA polymerase induced by growth feedback enhances overall effective cooperativity. Our results are consistent with this work as an increase in growth rate indeed expands the bistable range (Supplementary Fig. 6). In the case of the autoregulation of the T7 RNA polymerase, the circuit itself is not bistable but the nonlinear dilution included by the growth feedback increases the ultrasensitivity of the circuit and makes it bistable. In the case of the AraC self-activation circuit, it is bistable when growth feedback is decoupled or cell growth stops when the system reaches steady-state. However, under the conditions of constant growth rate with frequent dilution into the fresh medium, the activation threshold increases with the growth rate in an exponential way, making the activation of the switch infeasible in the biologically reasonable range of the inducer.

Thus, the effect of interactions between the circuit and the host is double-edged. While they endow new properties to some gene circuits, they impair or disguise the desired behaviors of others. Controlling strategies should be developed for the latter case. In this paper, we present a strategy by controlling the growth rate to minimize the effect of the growth feedback on the memory circuits. Instead of diluting the circuits in fresh medium, conditioned medium from the stationary phase or fresh medium with minimal nutrient can be used to slow down the growth rate and thus maintain the memory of the circuits.

### Conditioned medium from the stationary phase maintains the memory of the gene circuits

Conventionally, to test the functions of gene circuits, cells are often periodically diluted in fresh medium to keep them in the exponential phase. One of the underlying reasons for this is that the growth rate can be maintained at an almost constant value^7,23^. Another reason is that the primary metabolites are produced in the greatest quantities, which is important for the application of the synthetic gene circuits in industrial processes. While this dilution protocol worked successfully to demonstrate the properties of many circuits, it may cause some unexpected problems since the memory of the circuit can be lost, as shown here in this work. As during the stationary phase of bacteria, a surprisingly long period of constant protein production activity has been observed^41^. This is also confirmed in our work. The expression of the circuits can be maintained in the conditioned medium for at least 17 hours. Our results suggest that testing the gene circuit is feasible in the stationary phase, especially in the case where the growth is undesired. Given that wild bacteria in nature stay in the stationary phase most of their lifetime^41^, it is fascinating to test our synthetic circuits in this overlooked physiological state and to evaluate their functions when the growth-mediated feedback is decoupled. Of course, the components in the conditioned medium are very complex and may cause other problems. Future work is needed to evaluate and optimize this protocol.

### Bimodal distribution does not guarantee bistability

A bimodal distribution of gene expression with distinct ‘ON’ or ‘OFF’ subpopulation is usually taken as evidence of bistability of the system. However, there are several potential sources of bimodal distribution, including positive feedback loops in synthetic gene circuits^28,30^, ultrasensitivity and stochasticity^42,43^, the interplay between protein noise expression and network topologies^43^, the different growth rates of subpopulations^27,44,45^, and the feedback between cell growth and gene circuits^24,26^. To distinguish whether the designed circuit shows ultrasensitivity or bistability, hysteresis experiments are necessary to determine if the system can maintain its memory by reducing the concentration of the inducer. It is noted that the contribution of each source to bistability can be either additive or counteractive. Thus, it is interesting to quantify the contribution of each source to the generation of bistability. Here, the growth feedback is decoupled by controlling the cells in the non-growth conditions, and the contribution of the gene circuits to the bistability can be revealed.

### Resource relocation for the synthetic gene circuits complicates the circuit-host interactions

We demonstrated how the growth feedback affects the memory maintenance of two memory gene circuits and the underlying mechanism of how the self-activation switch is susceptible to, while the toggle switch is robust to the growth feedback. The primary factor is the perturbated concentrations of transcription activators and repressors due to cell growth, which might cause the activity changes of the circuits. However, the molecular mechanism of how gene activity changes during the fast cell growth phase remains unclear. One potential mechanism lies in the resource relocation strategy during cell division, as exogenous synthetic gene circuits have a lower priority to be expressed than the endogenous genes. That is, recourses such as ATP, as well as transcriptional and translational machinery components, are biased towards cell-cell-driven changes^46^. The resulting uncertainty needs to be controlled as well. One strategy is to integrate the synthetic gene circuits in the cell genome^47^. Another method is to manipulate the size of the resource pools^48,49^. Orthogonal ribosome pools or orthogonal DNA replication systems have been used to alleviate the effect of resource competition and gene coupling^50–52^. Insulation-based engineering strategies have shown to improve the resolution of the genetic circuit^53^. Future work is needed to further understand mechanisms of how circuit-host interactions affect the gene circuits and develop corresponding control strategies to design robust gene circuits.

## Material and Methods

### Strains, media and chemicals

E. coli DH10B (Invitrogen, USA) was used for all the cloning construction experiments. Measurement of the self-activated circuit was performed in E. coli K-12 MG1655ΔlacIΔaraCBAD as described in^13^. Measurement of the Toggle switch was performed with E. coli K-12 MG1655ΔlacI as described in^36^. Cells were grown in 5ml or 15ml tubes with 220 rotations per minutes at 37 °C in Luria-Bertani broth (LB broth) with 100μg/ml chloramphenicol or 50 μg/ml kanamycin. L-Arabinose (L-ara, Sigma-Aldrich) and anhydrotetracycline (aTc, Sigma-Aldrich) were dissolved in ddH2O and later diluted to appropriate working solution.

### Plasmids construction

The AraC self-activation circuit was constructed into either a pSB1C3 (high copy number, used for flow cytometry and plate reader analysis) or pSB3K3 (medium copy number, used for microfluidics) backbone according to the standard molecular cloning protocols using the standardized BioBricks parts from the iGEM Registry (www.parts.igem.org). The luxR self-activation circuit was constructed into pSB1C3 backbone. The araC gene was amplified by PCR using the BioBrick part BBa_C0080 as the template to have the lva-tag removed. The primers used were forward 5’-ctggaattcgcggccgcttctagatggctgaagcgcaaaatgatc-3’ and reverse 5’-ggactgcagcggccgctactagtagtttattatgacaacttgacggctacatc-3’. The BioBricks used were BBa_B0034 (ribosome binding site, RBS), BBa_K206000 (pBAD), BBa_K145015 (GFP with lva-tag), BBA_B0015 (transcriptional terminator) and BBa_C0062 (luxR). The sequence of pLux is 5’-acctgtaggatcgtacagggttacgcaagaaaatggtttgttatagtcgaataaa-3’. pLux was amplified by PCR using the BioBrick part BBa_R0062 as template. The primers used were forward 5’-gcttctagagacctgtaggatcgtacagggttacgcaagaaaatggtttgttatag-3’ and reverse 5’-ggactgcagcggccgctactagtatttattcgactataacaaaccattttc-3’. Detailed characterization of pLux can be found on website http://parts.igem.org/Part:BBa_R0062. All parts were first restriction digested using desired combinations of FastDigest restriction enzyme EcoRI, XbaI, SpeI and PstI (Thermo Fisher) and separated by gel electrophoresis, and then purified using GelElute Gel Extraction Kit (Sigma-Aldrich) followed by ligation using T4 DNA ligase (New England BioLabs). Then the ligation products were transformed into E. coli strain DH10B and later the positive colonies were screened. Finally, the plasmids were extracted using GenElute Plasmids Miniprep Kit (Sigma-Aldrich) and verified by sequencing. The operons constituting the self-activation circuits were constructed monocistronically. The plasmid of toggle switch was kindly provided by Dr. James Collins as described in^36,37^.

### Hysteresis experiment

#### Hysteresis experiment using conditioned medium

Cells were grown at 37°C on a shaker in 4ml LB broth using 15ml culture tubes. For the induction of self-activation, 2.5E-3% L-arabinose (Lara) were added to the growth medium at the beginning of the experiment. An identical culture without arabinose was grown in parallel. When the cells reached stationary phase, the cultures were centrifuged (2000g*5min) and supernatant from the parallel culture without L-ara was sterile-filtered and was used as the conditioned medium. For flow cytometry and plate reader analysis, the pellet from L-ara containing culture was first resuspended with the same volume of the conditioned medium as the volume of L-ara containing medium before the centrifuge; then the resuspended cells were diluted 100 folds into culture tubes containing 1ml conditioned medium with different concentrations of L-ara added. Finally, the cells were incubated in the shaker and measured at indicated time points for flow cytometry, or loaded onto 96-well plate for plate reader analysis. Hysteresis experiment using the conditioned medium for the toggle switch was performed the same way with the induction switched to aTc.

#### Hysteresis experiment using fresh medium

Cells were grown at 37°C in a shaker in 4ml LB broth using 15ml culture tubes. For the induction of self-activation, 2.5E-3% L-arabinose (L-ara) were added to the growth medium at the beginning of the experiment. When the cells reached the stationary phase, the cultures were centrifuged (2000g*5min) and the pellet was first resuspended with same volume of fresh medium without L-ara as the volume of L-ara containing the medium before the centrifuge; then the resuspended cells were diluted 100 folds into culture tubes containing 1ml fresh medium with different concentrations of L-ara added. Finally, the cells were incubated in the shaker and measured at indicated time points for flow cytometry, or loaded onto 96-well plate for plate reader analysis. Hysteresis experiment using fresh medium for the toggle switch was performed the same way with the induction switched to aTc.

### Flow cytometry

All samples were analyzed using Accuri C6 flow cytometer (Becton Dickinson) with excitation/emission filters 480nm/530nm (FL1-A) for GFP detection at indicated time points. 10,000 events were recorded for each sample. At least three replicated tests were performed for each experiment. Data files were analyzed with MATLAB (MathWorks).

### Dynamic analysis performed by Plate Reader

Synergy H1 Hybrid Reader from BioTek was used to perform the dynamic analysis. 200 μl of culture was loaded into each well of the 96-well plate. LB broth without cells was used as a blank. The plate was incubated at 37°C with orbital shaking at the frequency of 807cpm (circles per minute). OD (optical density) of the culture was measured by absorbance at 600nm; GFP was detected by excitation/emission at 485nm/515nm. All the measurements were taken at 30 minutes intervals.

### Microfluidics and microscopy

Cells carrying the circuit of self-activation or the toggle switch were grown in 5ml LB plus indicated induction and antibiotics overnight; then the cells were centrifuged at 2000g*5min followed by resuspension into 0.5 ml fresh medium with 0.75% Tween-80 (Sigma-Aldrich) and loaded onto the device. After the cells were loaded onto the device properly, fresh media with indicated induction was supplied to the cell for 16 hours; then the cells were switched to conditioned medium with indicated induction for various amounts of time as needed. Media switching was accomplished by adjusting the relative height of syringes containing fresh or conditioned medium. Details regarding design of the chip and setup of the device can be found in ref.^54^. Phase and green fluorescent images were taken every 15 min under the magnification of 40X with Nikon Eclipse Ti inverted microscope (Nikon, Japan) equipped with an LED-based Lumencor SOLA SE. Perfect focus was obtained automatically using Nikon Elements software.

## Mathematical modeling

### Self-activation Switch

The following mathematical model was used for the self-activation switch circuit without growth feedback,

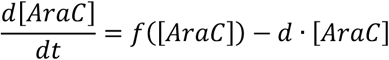

where 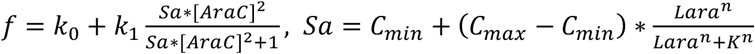, [AraC] is the concentration of AraC, which is co-expressed with GFP and thus used interchangeably. The derivation of the equation for the circuit can be found in our previous work^13^. The model is suited to analyze the steady-state behavior of the system under the no-growth condition or when the system reaches the stationary phase, where AraC steady-state level is controlled only by the production and degradation rate and depends on the circuit itself and thus shows the true behavior of the circuit. The hysteresis curves in Fig. 1B and the rate-balance plot in Fig.2E are based on this model.

In our system, the cell growth rate is dynamic in most cases as we diluted cells into the fresh medium at the beginning and cultured them overnight. Thus, the system covers all cell growth phases from exponential phase to stationary phase. To model the interplay between the cell growth and gene circuit, the mathematical model was revised by coupling cell growth.

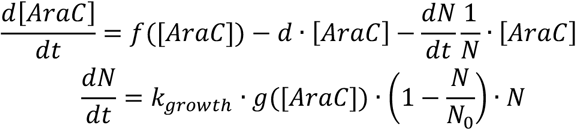

where *N* is the cell density, *k_growth_* is the maximum growth rate and *N*_0_ is the maximum cell density. The logistic growth function is used to describe dynamics of the cell density, in which the growth rate depends on the circuit expression 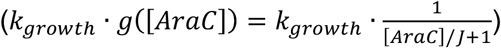. Fig. 2F is based on this revised model. When the system reaches the steady-state, cells growth stops and thus there are no changes in cell density 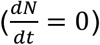, and as such, the third term in AraC equation is 0 and the system is the same as the one without growth feedback. That is, including the cell density equation does not affect the existence of bistability of the circuit but may change dynamics of the circuit. Fitted parameters are, unless otherwise mentioned, *C_min_* = 0.9, *C_max_* = 3, *n* = 3, *K* = 1.92 × 10^−3^, *k*_0_ = 0.07, *k*_1_ = 1.4, *d* = 0.7, *J* = 1, *k_growth_* = 2, *N*_0_ = 1.35, Lara = 0~2.5 × 10^−3^.

For the theoretical analysis, we also considered a scenario of constant growth rate to study how the strength of growth feedback affects the gene circuits. Constant growth rate can be achieved with frequent dilutions to control the cell in the exponential growth phase. In this case, the model is simplified as follows:

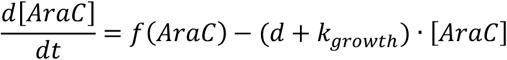

Supplementary Fig. 6 shows the simulation under several constant growth rates based on this model. With an increase in the growth rate constant, the steady-state level of AraC decreases and the activation threshold increases exponentially.

### Toggle Switch

The following equations were used the toggle switch circuit without growth feedback,

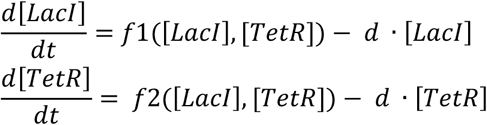

where 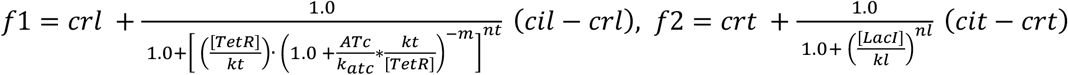, [*LacI*] and [*TetR*] are the concentration of LacI and TetR, respectively. LacI is co-expressed with GFP, and thus was used interchangeably. The derivation of the equation for the circuit can be found in our previous works^36,37^. The nullclines in Fig. 4J were based on this revised model.

The following mathematical model was used for the toggle switch circuit by coupling cell growth.

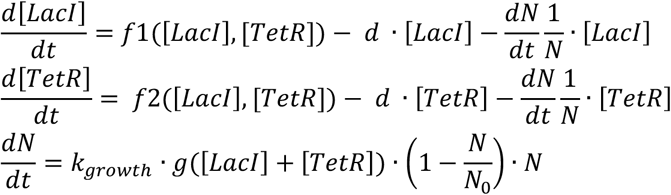

where 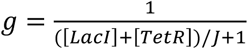. Supplementary Fig. 7 was based on this revised model. Fitted parameters are, unless otherwise mentioned, *crl* = 0.5, *crt* = 0.5, *cil* = 21.25, *cit* = 21.25, *d* = 0.5, *m* = 3, *nl* = 3.5, *nt* = 1.5, *kl* = 8, *kt* = 6, *k_atc_* = 0.4, *J* = 20, *k_growth_* = 2.3, *N*_0_ = 1.35, *ATc* = 0~4.

### Stochastic simulation of cell division events at the single-cell level

To simulate the interplay between the circuit and cell growth at the single-cell level by considering the stochasticity from cell division before the system reaches the stationary phase, we developed the following algorithm.

1. Initialize the system: cell number *N*, time *t* = 0, gene expression profile *x_i_* for each cell.
2. Increase time by a small step *t* = *t* + Δ*t*.
3. Calculate the probability of division for each cell *P_i_* according to

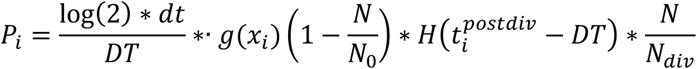

where *DT* is the cell doubling time, 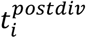 is the time after last division for each cell, *N_div_* is the number of ready-to-division cells based on the condition *t_postdiv_* > *DT*, *H* is the Heaviside step function, *g*(*x*) indicates on the effects of circuit expression on the growth rate.
4. Generate uniformly distributed random number *r_i_* between [0 1] for each cell. If *r_i_* > *P_i_*, the cell divides. Update 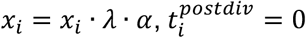, add a new cell *x_new_* = *x_i_* · *λ* · (1 – *α*), 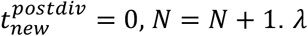 represents the fold change of *x_i_* (0.5~0.7) for each cell division by considering the expression changes before the cell division. *α* is used for simulation of unsymmetrical cell division. Otherwise, 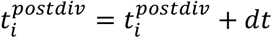.
5. Simulate the dynamics of gene circuit for each cell with the current *x_i_* according to the ODEs of the circuit without growth feedback.
6. Repeat 2~5 until *t* is more than maximum time *T_max_*.

The effects of the cell size on the concentration of variables were not considered in the model. Fig. 2E and Fig. 4J, four cell divisions at 0.7h, 1.4h, 2.1h 3.15h were used as one example for demonstration and comparison in both the self-activation switch and the toggle switch circuits.

## Data availability

The data that support the findings of this study are available on request from the corresponding author.

## Code availability

The modeling code for deterministic and stochastic numerical simulations is available on request from the corresponding author.

## Author Contributions

X-J.T. conceived the study. X-J.T., R.Z., J.L., Q.Z. and X.W. designed the study. R.Z., J.L., P.S. and X.C. performed experiments. J.M., H.G and X-J.T. performed model studies. R.Z., X-J.T., and X.W. analyzed the data and wrote the manuscript with input from all authors.

## Acknowledgments

We thank X FU, T. Hong, G. Yao, J Xing, and T Hwa for valuable comments. This project was supported by the ASU School of Biological and Health Systems Engineering and NSF grant (EF-1921412) (to X-J.T.), and NIH grant (GM106081) (to X.W.). H Goetz and J Melendez-Alvarez were also supported by the Arizona State University Dean’s Fellowship.

