## Supplementary Figures 1-9 for "Topology-Dependent Interference of Circuit Function by Growth Feedback"

This PDF file includes:
Supplementary Figures 1-9
Supplementary Video 1-2

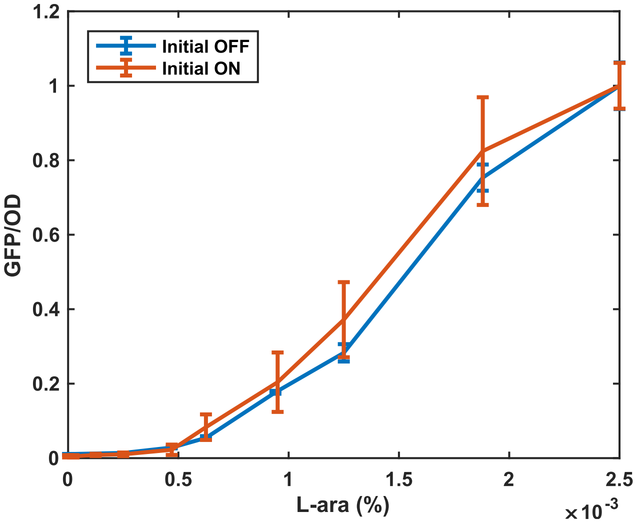


**Supplementary Figure 1. Similar dose-response curves with different initial states.** The steady-state level of GFP/OD as a function of the L-ara concentration with the initial state “ON” (red curve) and ‘OFF (blue curve). Mean value of three replicates were shown.

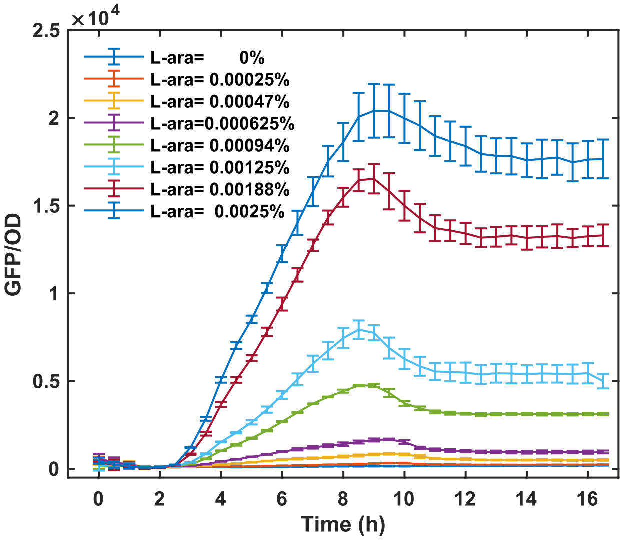


**Supplementary Figure 2.** Dynamics of GFP/OD after 1:100 dilution of 'OFF' cells into fresh medium with various concentrations of L-ara. Mean value of three replicates were shown.

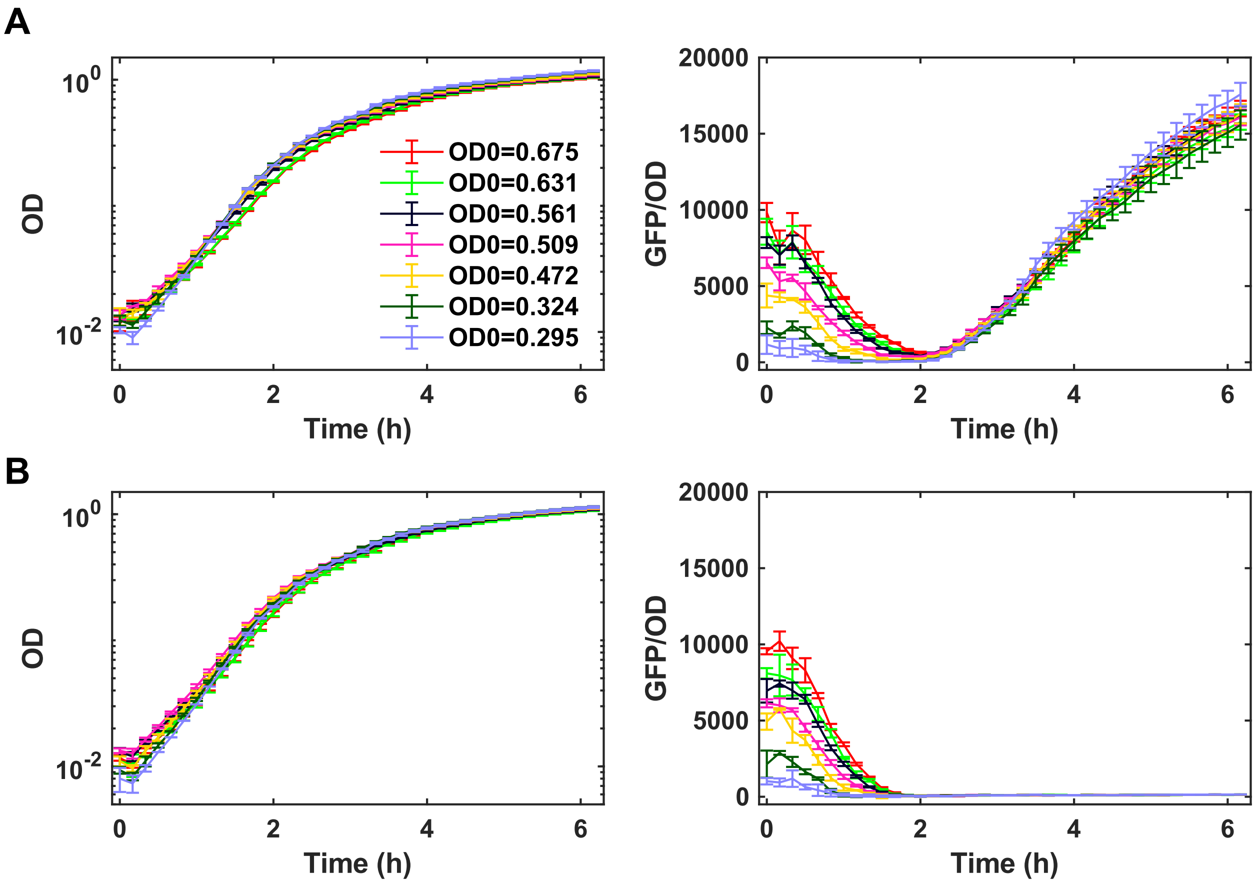


**Supplementary Figure 3. Memory loss of self-activation circuit was confirmed with dilution of cells in the exponential phase into fresh medium.** The dynamics of cell growth (OD) and GFP after dilution of cell that were activated to different levels at OD=0.3~0.7 as indicated into fresh medium with high dose of ($2.5*10^{-3}$%) (A) or no L-ara (B). The OD before dilution was shown as indicated.

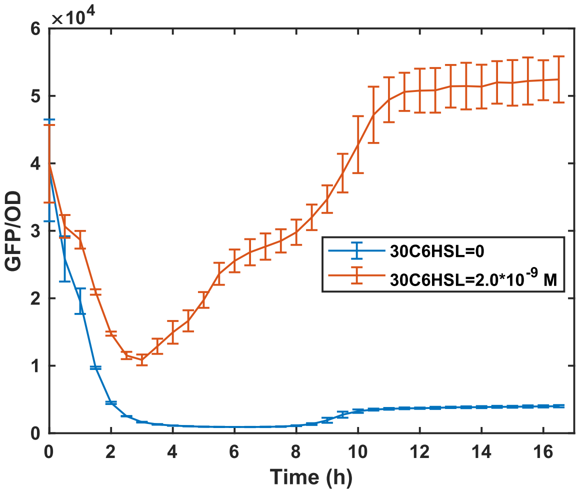


**Supplementary Figure 4. Loss of memory of the LuxR self-activation circuit.** The dynamics of GFP after dilution of activated cells into fresh medium with high dose or no inducer 30C6HSL.

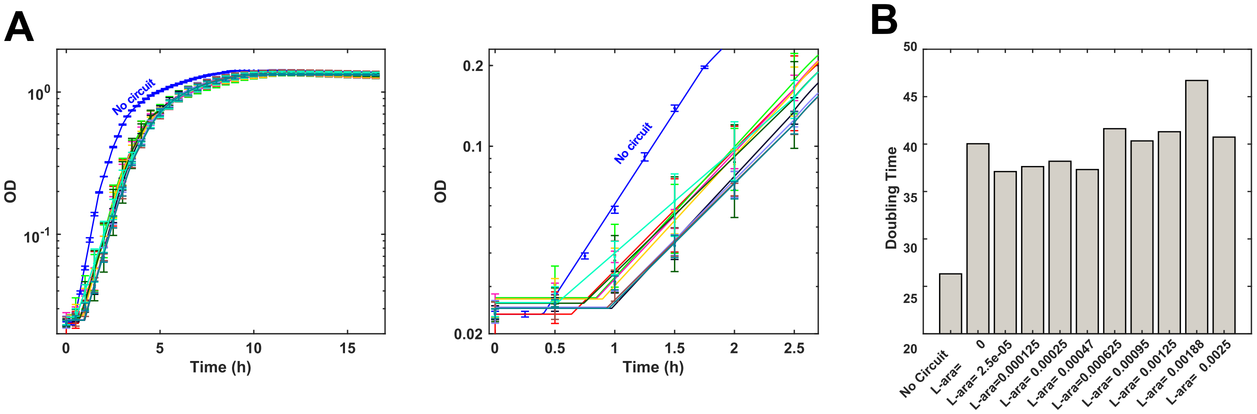


**Supplementary Figure 5. Inhibition of cell growth by gene circuit expression.**

(A) The growth curve without or with activation of the self-activation circuit.

(B) The calculated doubling time in the exponential growth phase based on the fitting growth curve.

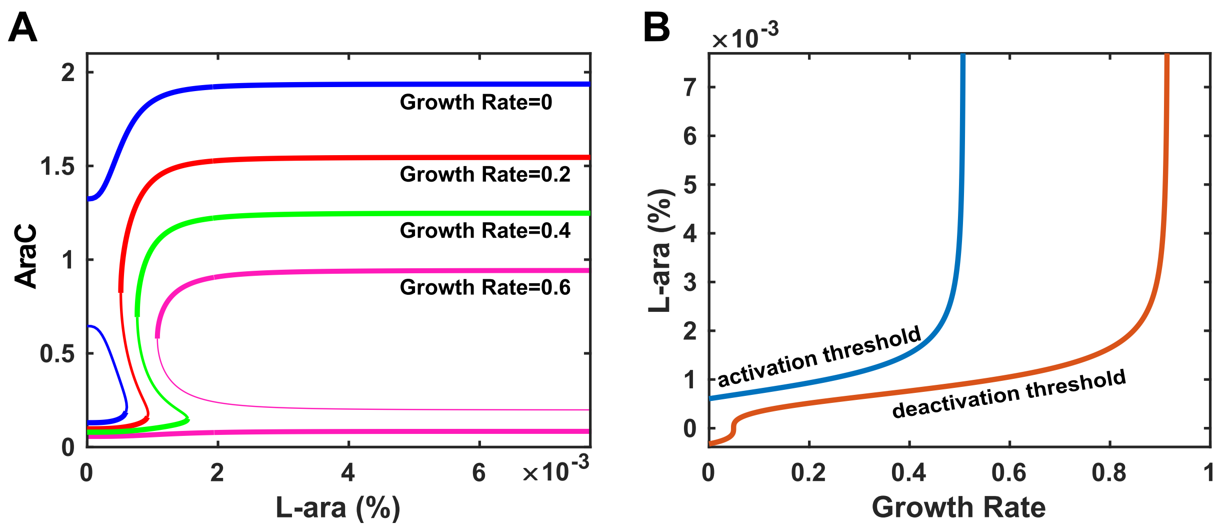


**Supplementary Figure 6. The effects of growth rate on the bistability of the self-activation circuit.**

(A) Bifurcation diagram of AraC versus the L-ara level under conditions with various constant growth rates.

(B) Dependencies of the thresholds for activation and deactivation of the circuit on growth rate. Under the conditions with constant growth rate, the activation threshold of L-ara increases with growth rate exponentially.

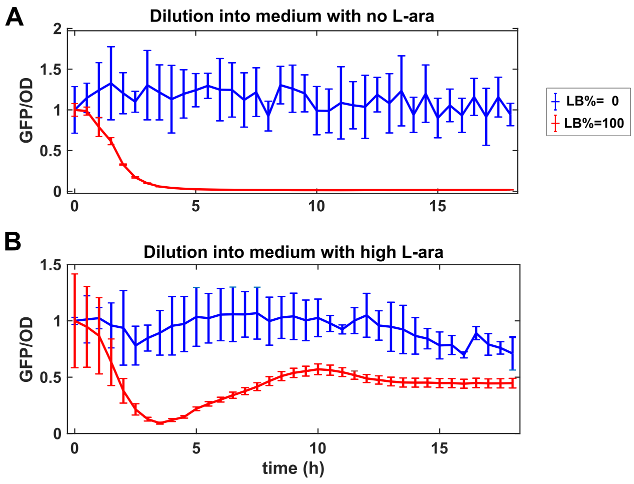


**Supplementary Figure 7. Decoupling of growth-mediated feedback with low-nutrient medium maintains the memory of the self-activation circuits.**

The dynamics of the GFP/OD in the self-activation circuit after diluting activated cells into mediums without (A) or with high Lara (B), and low-nutrient (blue lines) or rich-nutrient (red lines).

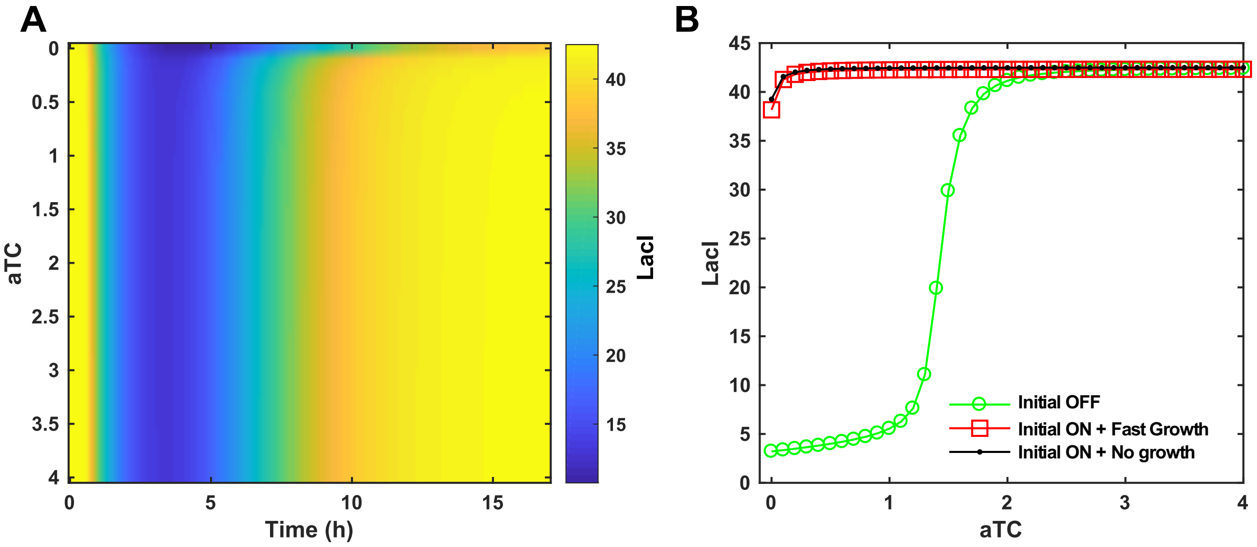


**Supplementary Figure 8. Simulation confirms that the toggle switch is refractory to growth feedback.**

**(A)** Simulation with mathematical model shows the change of LacI as a function of time and dose of inducer aTc. The system was set to the activated (LacI high) state initially.

**(B)** The hysteresis curve for the toggle switch circuit coupled with and without growth feedback.

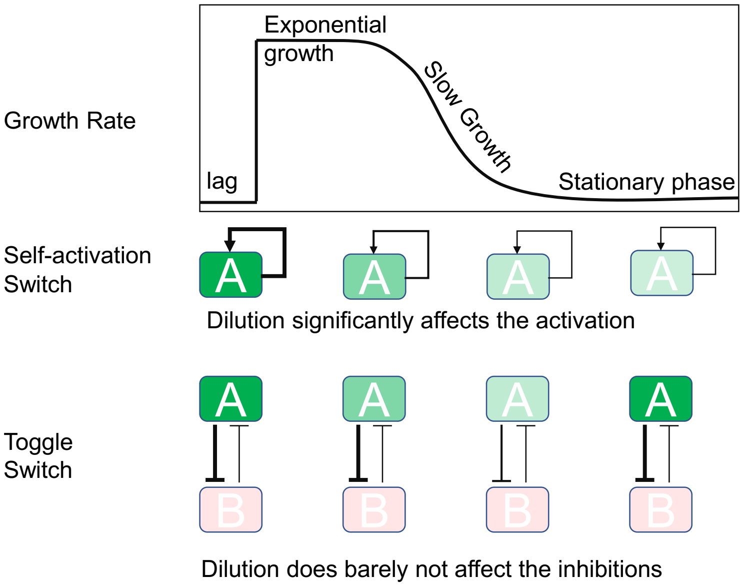


**Supplementary Figure 9. Schematics of how self-activation switch and toggle switch behave differently during the process of cell growth after cells dilution into fresh medium.** Dilution of the gene expression significantly affects the production rate of the gene in self-activation switch and thus leads the inactivation of the switch. However, in the toggle switch, dilution of the gene expression does not affect the production rate much and relative expression levels of two mutual-inhibitive genes, and thus lead to the robust memory.

**Supplementary Video 1**. A time-lapse movie showing the dynamics of GFP in the AraC self-activation circuit under the condition without L-ara fresh medium for 14h and conditioned medium for 7 hours thereafter. Initially the system was set in the high GFP state. The photos were taken at 15 minutes intervals.

**Supplementary Video 2**. A time-lapse movie showing the dynamics of GFP in the AraC self-activation circuit under condition with a high dose of L-ara and fresh medium condition for 14h and conditioned medium for 7 hours thereafter. Initially the system was set in the high GFP state. The photos were taken at 15 minutes intervals.

**Supplementary Video 3**. A time-lapse movie showing the dynamics of GFP in the toggle switch circuit under the condition without aTc and fresh medium for 16h and conditioned medium for 10 hours thereafter. Initially the system was set in the high GFP state. The photos were taken at 15 minutes intervals.

**Supplementary Video 4**. A time-lapse movie showing the dynamics of GFP in the toggle switch circuit under the condition with 2ng/ml aTc and fresh medium for 16h and conditioned medium for 10 hours thereafter. Initially the system was set in the high GFP state. The photos were taken at 15 minutes intervals.
